# The self-organization model reveals systematic characteristics of aging

**DOI:** 10.1101/502815

**Authors:** Yin Wang, Tao Huang, Xianzheng Sha, Yixue Li, Chengzhong Xing

## Abstract

Aging is a fundamental biological process, where key bio-markers interact with each other and synergistically regulate the aging process. Thus aging dysfunction will induce many disorders. Finding aging markers and re-constructing networks based on multi-omics data (i.e. methylation, transcriptional and so on) are informative to study the aging process. However, optimizing the model to predict aging have not been performed systemically, although it is critical to identify potential molecular mechanism of aging relative diseases.

This paper aims to model the aging self-organization system using a serious of supervised learning methods, and study complex molecular mechanism of aging at system level: i.e. optimizing the aging network; summarizing interactions between aging markers; accumulating patterns of aging markers within module; finding order-parameters of the aging self-organization system.

In this work, the normal aging process is modeled based on multi-omics profiles across tissues. In addition, the computational pipeline aims to model aging self-organizing systems and study the relationship between aging and related diseases (i.e. cancers), thus provide useful indexes of aging related diseases and improve diagnostic effects for both pre- and pro- gnosis.

## Introduction

Aging is a complex process regulated by key bio-markers, reflecting disorders / declined abilities of tissues [1]. Dysfunction of aging has been shown to relate many diseases, such as diabetes, Parkinson disease [2], Alzheimer’s disease [3] and cancers [4]. As a result, finding aging markers is critical to study aging related diseases and identify healthy genomic diagnostics (i.e. by predicting the chronological age (group) based on molecular profiles). For example, multi-tissue predictors of age have been calculated by methylation [5] or mRNA expression profiles[6]; and there are more age predictors based on single tissue (e.g. brain [7], breast [8], ans so on), also provide insights on aging related diseases [6] (i.e. cancers, Alzheimer’s disease).

Further, the aging markers interact with each other [9], and synergetically coordinate the aging process, herein generating the self-organization system [10] of aging, where particular bio-markers regulate the aging process in different age groups, respectively. Although tissues become disordered / functional decline during aging in general (often evaluated by the entropy [11]), a serious of aging markers perform special / ordered functions in special aging stages / age groups. In addition, these markers / genes interact with each other, coordinately involving in the aging process; therefore, the interactions / modules between such markers also provide critical patterns of aging. In summary, finding bio-markers to predict the chronological ages, summarizing interactions between aging markers, and optimizing the aging self-organization system based on molecular profiles (i.e. methylation, expression and so on) of normal tissues from healthy persons, could help predict future health risks at system level. However, these works have not been solved entirely for the aging process.

In this work, we modeled the aging self-organization system using a serious of computational methods: filtering inter-connection networks between different age groups by the maximum mutual information and minimum redundancy criterion in the information theory; summarizing interactions between bio-markers by the convolution technology; calculating patterns by accumulating weighted genes within the same module; selecting module scales by the hierarchical clustering method and cross validation; identifying order parameters in the aging self-organization system by network sparsification.

The prediction results showed high classification accuracy between different age groups; moreover, the enrichment analysis and network analysis also found key functions of the order parameters. Thus critical complex characteristics (i.e. hierarchies, emergencies and bifurcations) were identified in different aging stages. Aging acceleration patterns were also identified across cancers. In short, the aging process can be thorough studied by modelling the aging self-organization system.

## Results and discussion

### A brief description of the aging self-organization system

In the aging self-organization system, genes interacted with each other, and synergetically coordinated the aging process. Therefore, the aging process could also be evaluated by interactions between aging markers. The aging markers clustered nearby would drive similar function [12], and could be summarized within the same module. Each module take a particular part during aging, and regulated the aging process altogether. Different levels / hierarchies of the markers / modules reflected special the complexity of the aging system, herein reflecting particular patterns in different aging stages. As a result, the aging self-organization system would emerge important characteristics apart from any single isolated marker / module. In summary, modules based on aging markers coordinately determined the bifurcations and displayed critical differential patterns between age groups, where key aging markers within modules could be identified as the order parameters in the aging self-organization system.

Further, the system from pathological samples should be deviated the normal aging self-organization system (from healthy persons): for example, the aging system with disease (i.e. cancers) should display significant acceleration from the normal aging process. Accordingly, the following parts of this paper depicted results of modelling the aging self-organization system and the computational pipeline was shown in Figure 1.

**Figure 1.**
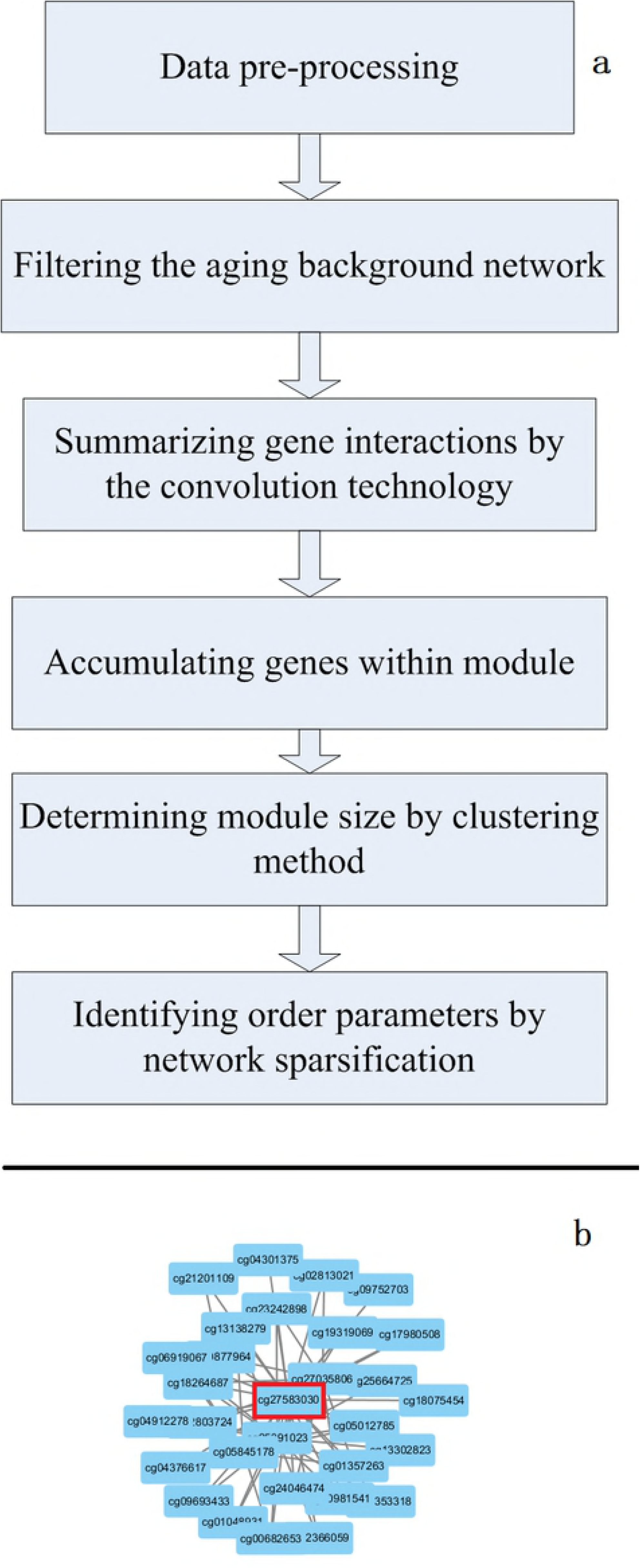
Overview of the aging self-organization system. (a) the computational pipeline of modelling the aging self-organization system; (b) an example of the convolution of interactions between methylation cg27583030 (SLC25A4) and other genes in the model of age group 50-70 vs. 70-survival;

### Classification results of the aging self-organization system

The expression and methylation profiles were used to test the classification abilities of the aging self-organization system between different age groups, respectively (Figure 2 and 3). Table 1–3 showed that classification results based on the self-organization system have lower error rates than traditional feature selection methods (i.e. the refieff-mRMR pipeline [13]), using both methylation and expression data between age groups. As a result, the self-organization system reduced the feature dimensions effectively and extracted critical modules by finding the order-parameters, and identifying key differences based on aging markers / interactions in the aging process at system level.

**Figure 2.**
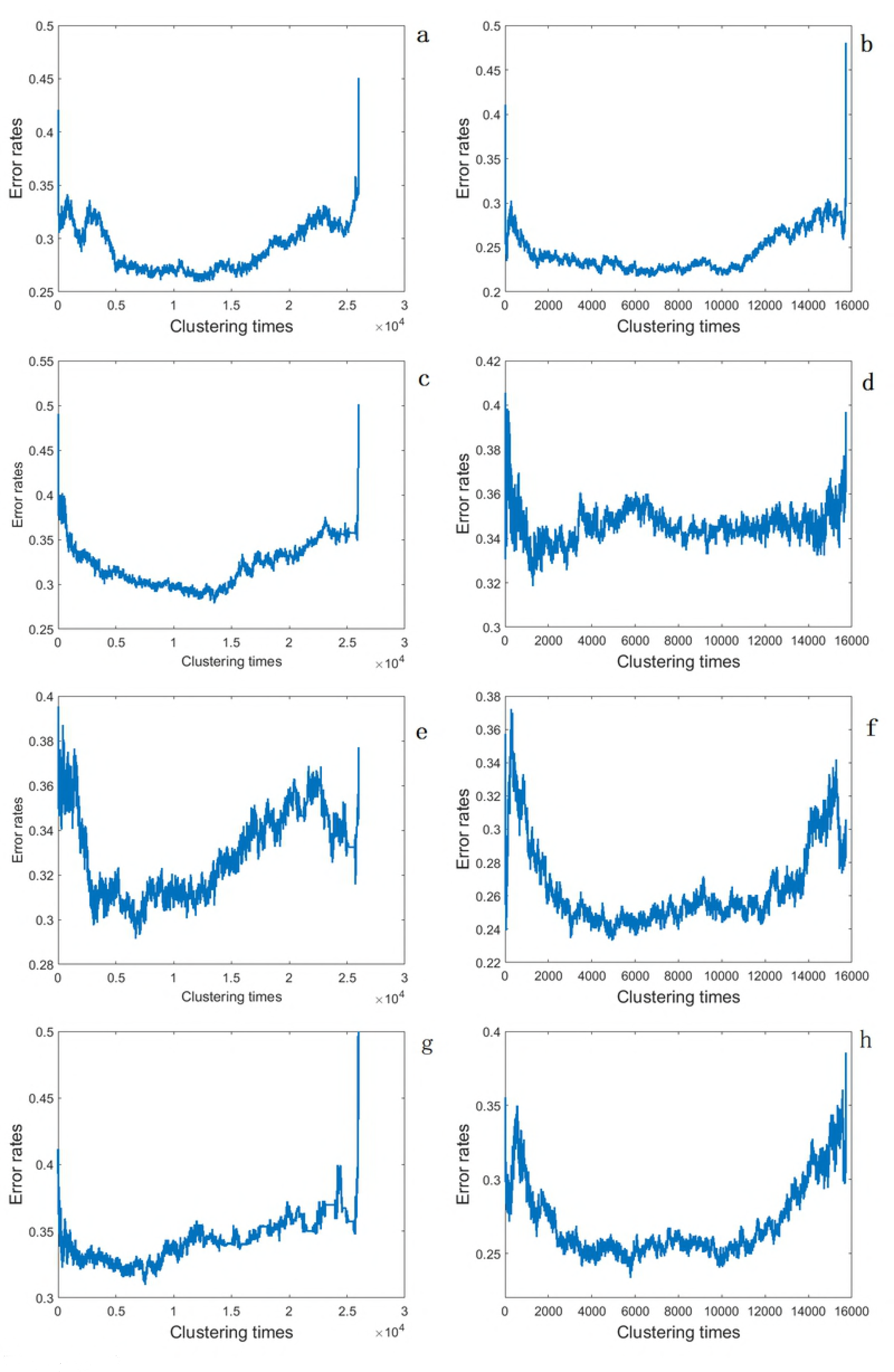
Learning curves of the the aging self-organization system. (a, c, e, g) methylation profiles; (b, d, f, h) expression profiles; (a, b) 0-50 vs. 50-survival; (c, d) 0-20 vs. 20-50; (e, f) 20-50 vs. 50-70; (g, h) 50-70 vs. 70-survival;

**Figure 3.**
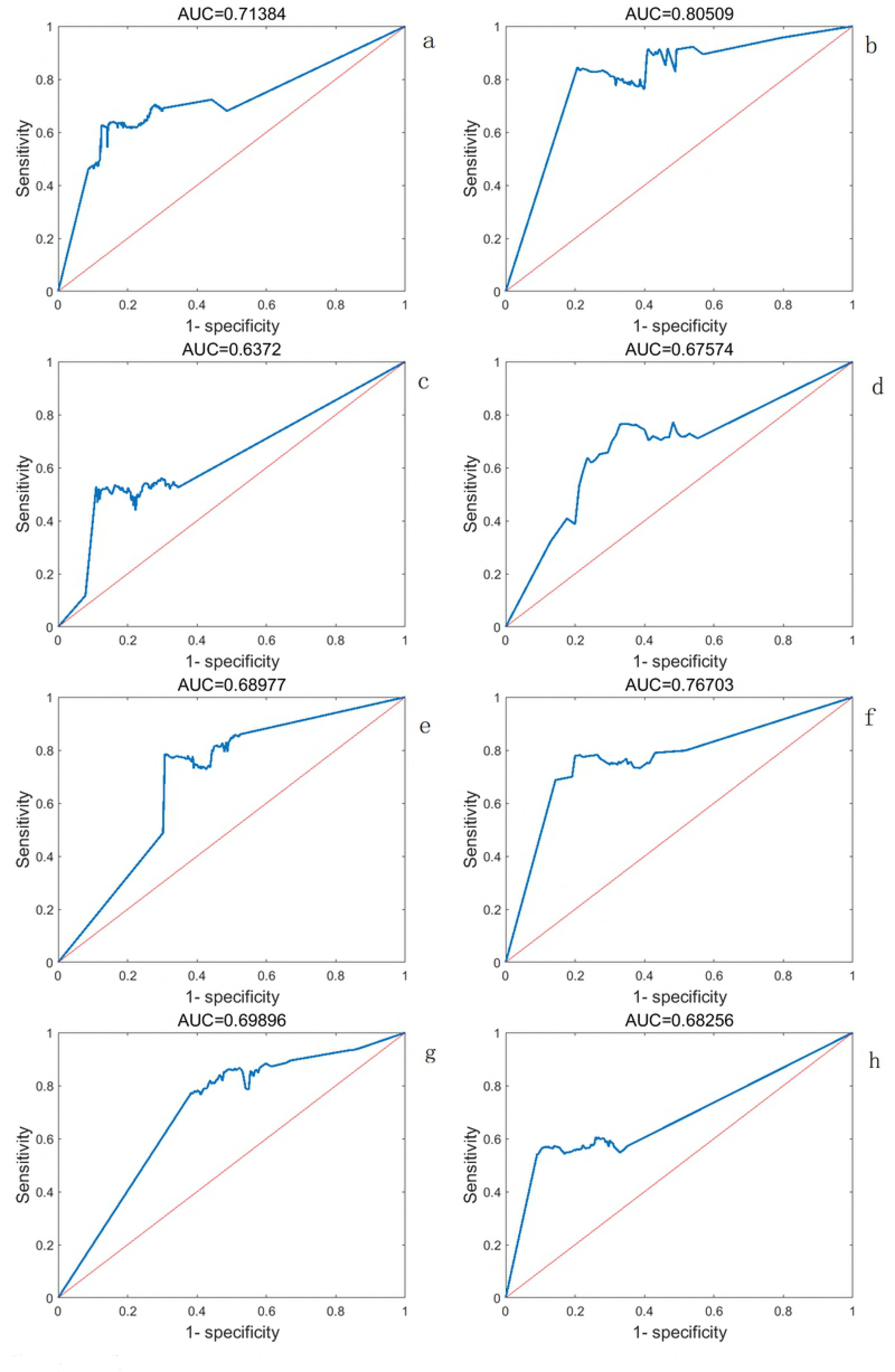
ROC curves of the the aging self-organization system in test data. (a, c, e, g) methylation profiles; (b, d, f, h) expression profiles; (a, b) 0-50 vs. 50-survival; (c, d) 0-20 vs. 20-50; (e, f) 20-50 vs. 50-70; (g, h) 50-70 vs. 70-survival;

**Table 1.** overview of the aging self-organization system

| model | profiles | Samples<br>(training data + test<br>data) | Order parameters<br>(original profiles +<br>summarized<br>interactions) | Modules (dimensions) |
| --- | --- | --- | --- | --- |
| 0-20 vs.<br>20-50 | methylation | 1760+798 | 4360+6408 | 2683 |
| 20-50 vs.<br>50-70 | methylation | 1875+845 | 2137+6713 | 2920 |
| 50-70 vs.<br>70-survival | methylation | 1285+585 | 1484+5497 | 1918 |
| 0-50 vs.<br>50-survival | methylation | 3045+1381 | 4825+4891 | 2746 |
| 0-20 vs.<br>20-50 | expression | 655+315 | 246+2211 | 1173 |
| 20-50 vs.<br>50-70 | expression | 975+473 | 2658+4660 | 2388 |
| 50-70 vs.<br>70-survival | expression | 845+411 | 2201+6044 | 2454 |
| 0-50 vs.<br>50-survival | expression | 1500+726 | 2832+4413 | 2067 |

**Table 2.**
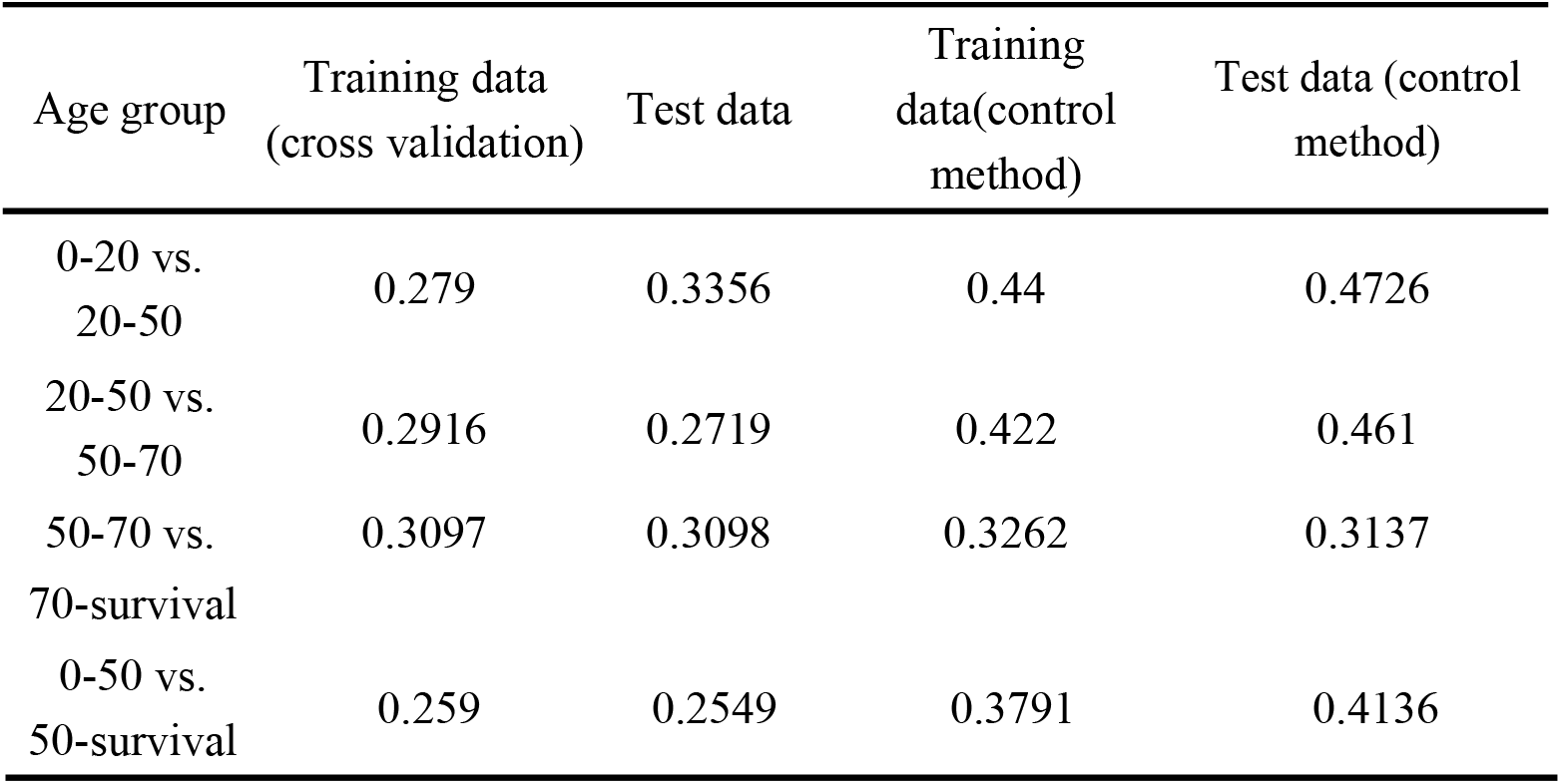
classification results (error rates) based on methylation data

**Table 3.** classification results (error rates) based on expression data

| Age group | Training data<br>(cross validation) | Test data | Training data<br>(control method) | Test data (control<br>method) |
| --- | --- | --- | --- | --- |
| 0-20 vs.<br>20-50 | 0.3185 | 0.3084 | 0.351 | 0.3911 |
| 20-50 vs.<br>50-70 | 0.2333 | 0.2308 | 0.2613 | 0.4325 |
| 50-70 vs.<br>70-survival | 0.2337 | 0.279 | 0.2726 | 0.4659 |
| 0-50 vs.<br>50-survival | 0.2161 | 0.2066 | 0.3492 | 0.4297 |

### Biological features of the order parameters

The methylation order parameter with the maximum relieff weight was the convolution interactions of cg27583030 (SLC25A4, weight=0.1974, shown in Figure 1b) in the model of age group 50-70 vs. 70-survival. Common SLC25A4 related pathway were apoptosis and survival regulation of apoptosis by mitochondrial protein, reflecting the relationship between aging and cellular apoptosis [14]. The expression order parameters with the largest relieff weight was original profile of NRBF2 (weight=0.2493) between age group 50-70 vs. 70-sruvival. NRBF2 played a role in cellular survival and neural progenitor cell survival during differentiation [15], and dysfunction of NRBF2 also affected the aging process.

Further, enrichment analyses of the order parameters in each module were performed on Biological Process (BP) terms of Gene Ontology (GO) and KEGG pathways using the hypergeometric test (Table 4–5 and S1-S2). The most significant BP term was the negative regulation of viral process (GO:0048525, fdr=0.0005) in the 245th module of the model between age group 20-50 vs. 50-70 based on the methylation profiles, reflecting the relationship between the immunity system and aging [16]; and the most significant KEGG pathway was Phenylalanine metabolism (fdr=0.0009) based on the methylation profiles in the model between age group 0-50 vs. 50-survival, indicating the critical metabolism during aging. In addition, the annotation of order-parameters also reveal key functions across different aging stages, i.e. BP terms were enriched in aging related diseases in the early stage of aging, and enriched in tissue dysfunction in the later stage. It is perhaps functional decline of the immunity system induced aging / tissue dysfunction.

**Table 4.**
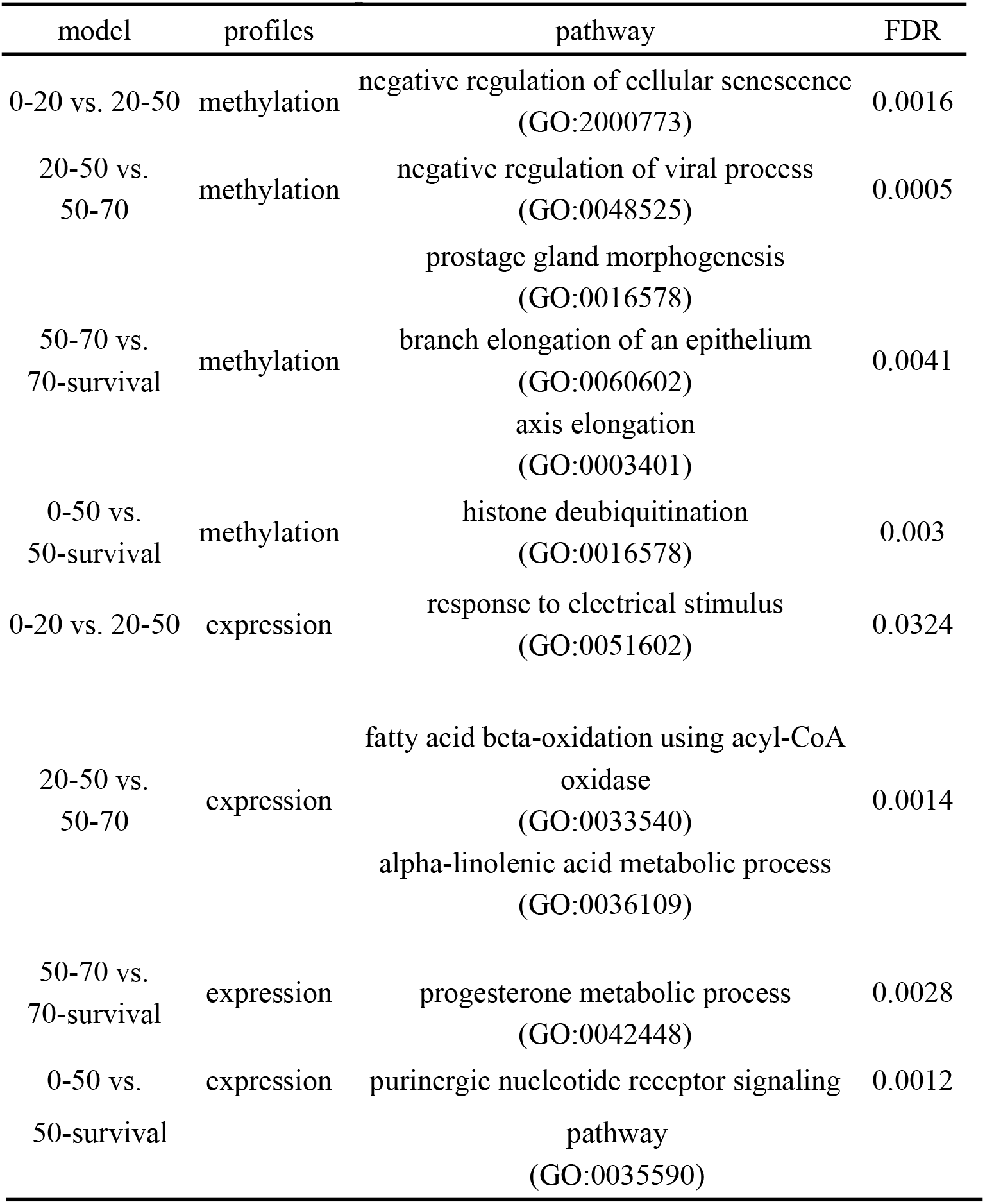
top enriched BP terms within module

**Table 5.** top enriched KEGG pathways within module

| model | profiles | pathway | FDR |
| --- | --- | --- | --- |
| 0-20 vs.<br>20-50 | methylation | Maturity onset diabetes of the young | 0.0019 |
| 20-50 vs.<br>50-70 | methylation | Focal adhesion | 0.0015 |
| 50-70 vs.<br>70-survival | methylation | Homologous recombination | 0.0014 |
| 0-50 vs.<br>50-survival | methylation | Phenylalanine metabolism | 0.0009 |
| 0-20 vs.<br>20-50 | expression | Lysosome | 0.0104 |
| 20-50 vs.<br>50-70 | expression | Retinol metabolism | 0.0022 |
| 50-70 vs.<br>70-survival | expression | Prion diseases | 0.0027 |
| 0-50 vs.<br>50-survival | expression | Long-term potentiation | 0.0035 |

Strikingly, enriched functions across modules indicated the common themes of aging (Table 6 and 7). For example, BP terms of organ morphogenesis were enriched during both young (0-20 vs. 20-50) and old (50-70 vs. 70-survival) age groups, reflecting the basal role of tissues affected by the aging process. Moreover, cancer and related signaling KEGG pathways were enriched across different aging stages based on both methylation and expression profiles (Figure 4). These results reflected the cross-talk between aging and cancer, where dysfunction of aging might indicate diseases / cancerization of tissues.

**Table 6.**
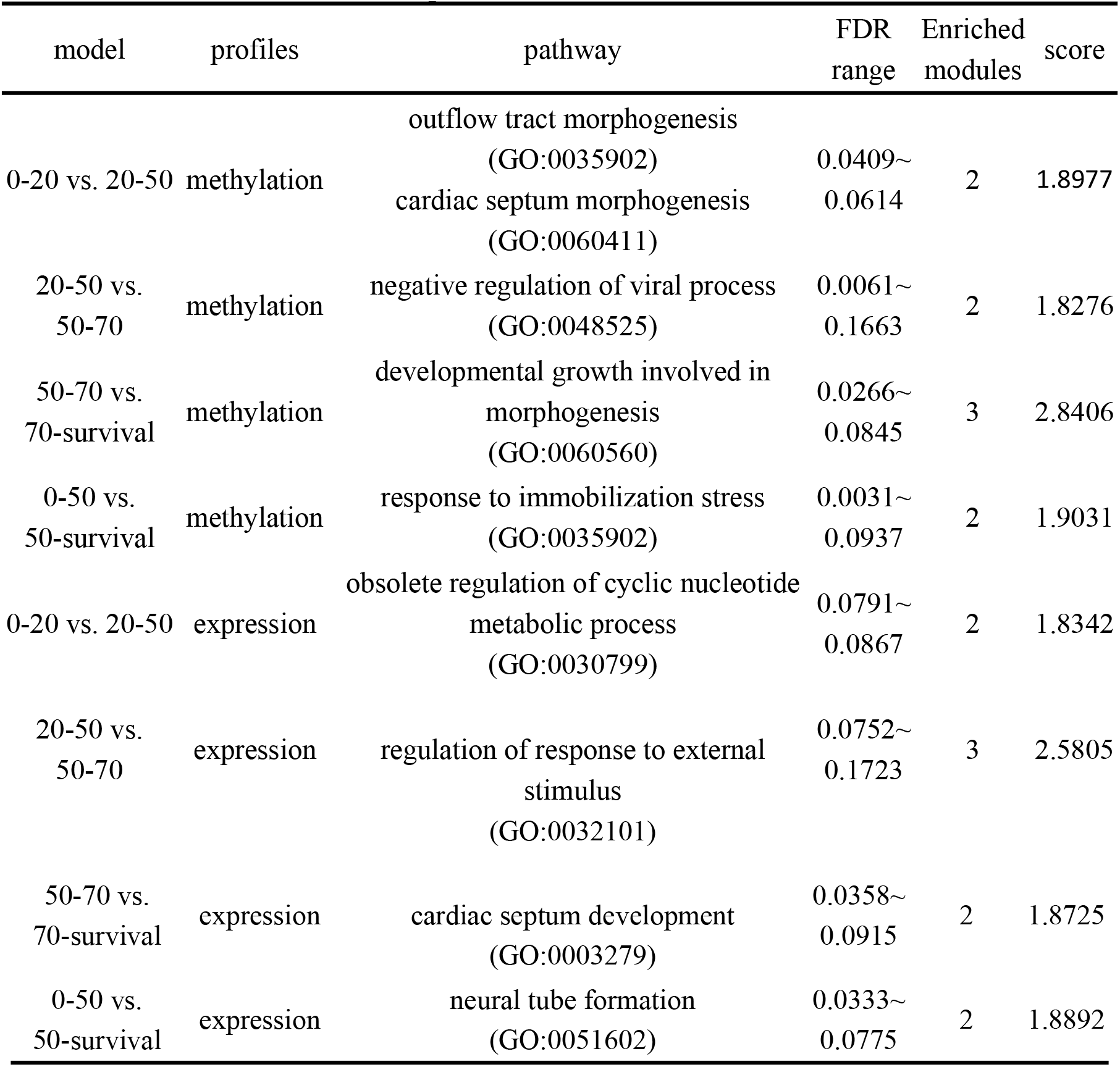
top enriched BP terms across modules

**Table 7.** top enriched KEGG pathways across modules

| model | profiles | pathway | FDR range | Enriched<br>modules | score |
| --- | --- | --- | --- | --- | --- |
| 0-20 vs.<br>20-50 | methylation | Glioma | 0.0193~0.184 | 11 | 9.8185 |
| 20-50 vs.<br>50-70 | methylation | Prostage cancer | 0.0922~0.185 | 16 | 13.6894 |
| 50-70 vs.<br>70-survival | methylation | Pancreatic cancer | 0.0714~0.1931 | 9 | 7.9217 |
| 0-50 vs.<br>50-survival | methylation | Pancreatic cancer | 0.041~0.1988 | 17 | 14.7595 |
| 0-20 vs.<br>20-50 | expression | ERBB signaling pathway<br>VEGF signaling pathway<br>Non-small cell lung cancer | 0.0902~0.1844 | 5 | 4.3889 |
| 20-50 vs.<br>50-70 | expression | Insulin signaling pathway | 0.0213~0.1858 | 10 | 8.8731 |
| 50-70 vs.<br>70-survival | expression | VEGF signaling pathway<br>Fc epsilon RI signaling pathway | 0.0134~0.194 | 9 | 7.8764 |
| 0-50 vs.<br>50-survival | expression | Prostage cancer | 0.0692~0.1887 | 11 | 9.4625 |

**Figure 4.**
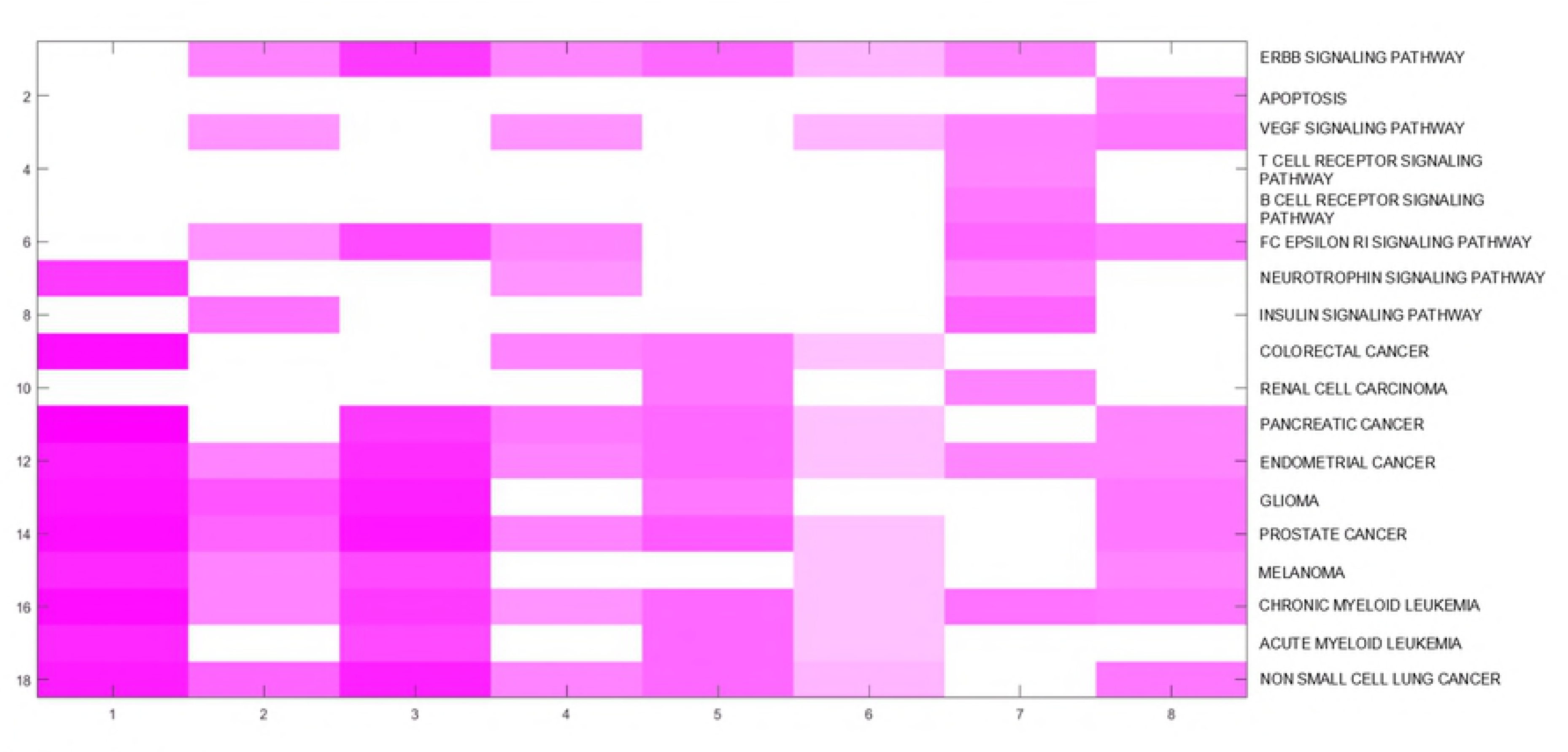
Biological features of the the aging self-organization system. The top 10 enriched KEGG pathways across different models of aging self-organization systems.

### Complexity characteristics of the aging self-organization system

In the self-organization system, molecules interacted with each other, and synergetically regulate particular process (i.e. aging). For example, summarized interactions of both methylation cg27583030 (SLC25A4) (with higher relieff values, shown in Figure 1b). Moreover, a serious of order parameters were with summarized interactions other than the ordinary profiles, indicating functions of cross-talks between key markers during aging (Table 1).

In addition, any single isolated marker / module could not reflect all of the key characteristics between age groups effectively, but combination of modules could. It should be that cross-talks across modules generated the aging self-organization system with key differential patterns / bifurcations between age groups. Thus the hierarchies across modules promoted emergencies of the aging self-organization system, which were not displayed by any single module / order parameter. Therefore, the correspondences across modules indicated the hierarchies / emergencies of the aging self-organization system.

The corresponding networks across modules (also using the maximum mutual information minimum redundancy criterion) indicated core modules in the hierarchies of the self-organization system (Figure S1). Based on the methylation profiles, the 492th module were with the maximum interaction score (mean value=0.0189), connecting other 2682 modules (in the model of age group: 0-20 vs. 20-50). This module acted as a “hub” and connected 45 key BP functions (i.e. negative regulation of cellular senescence, regulation of attachment of spindle microtubules to kinetochore and negative regulation of potassium ion transmembrane transporter activity) and 60 KEGG pathways (i.e. Maturity onset diabetes of the young and Pentose and glucuronate interconversions), reflecting the crosstalk between immunity and key metabolic pathway during aging. Based on the expression profiles, the 1799th module were with the maximum interaction score (mean value=0.0762), connecting other 2387 modules (in the model of age group: 20-50 vs. 50-70). The module connected 46 key BP functions (i.e. fatty acid beta oxidation using acyl coa oxidase, cell aggregation and cytokine production) and 51 KEGG pathways (i.e. Retinol metabolism, alpha linolenic acid metabolism and Pyruvate metabolism), revealed basal metabolism pathways during aging. In short, the corresponding networks reflected correlations of important parts / functions in the aging self-organization system, such as the immunity system, cancer related pathways, and so on. It might be the cross-talk between the immunity system and cancers affected emergencies of critical themes in the aging process.

Therefore, the bifurcations (differential patterns) between age groups were investigated, where order parameters with convoluted interactions were summarized by each relieff weight, indicating order-disorder patterns of interactions between aging order-parameters from low (near the ordered / same pattern with low entropic values of the aging system) to high (near disordered / different patterns with high entropic values) values, or vise verse. As a result, significant differential patterns were found between age groups using both methylation and expression profiles (Figure 5). In the early aging stage (0-20 vs. 20-50), the aging self-organization systems were with significantly ordered patterns based on both methylation and expression profiles. As aging was regulated by special markers / order-parameters, the self-organization system showed ordered patterns in the early aging stage. However, the self-organization systems were with disordered patterns in middle (20-50 vs. 50-70) and later (50-70 vs. 70-survival) stage based on expression and methylation profiles, respectively, indicating tissue declined function / disordered patterns during aging. The aging process was driven by both special markers / order parameters (with order patterns) and tissue declined function (with disordered patterns), perhaps determined by the former in the early aging stage, and by the latter in the later stage.

**Figure 5.**
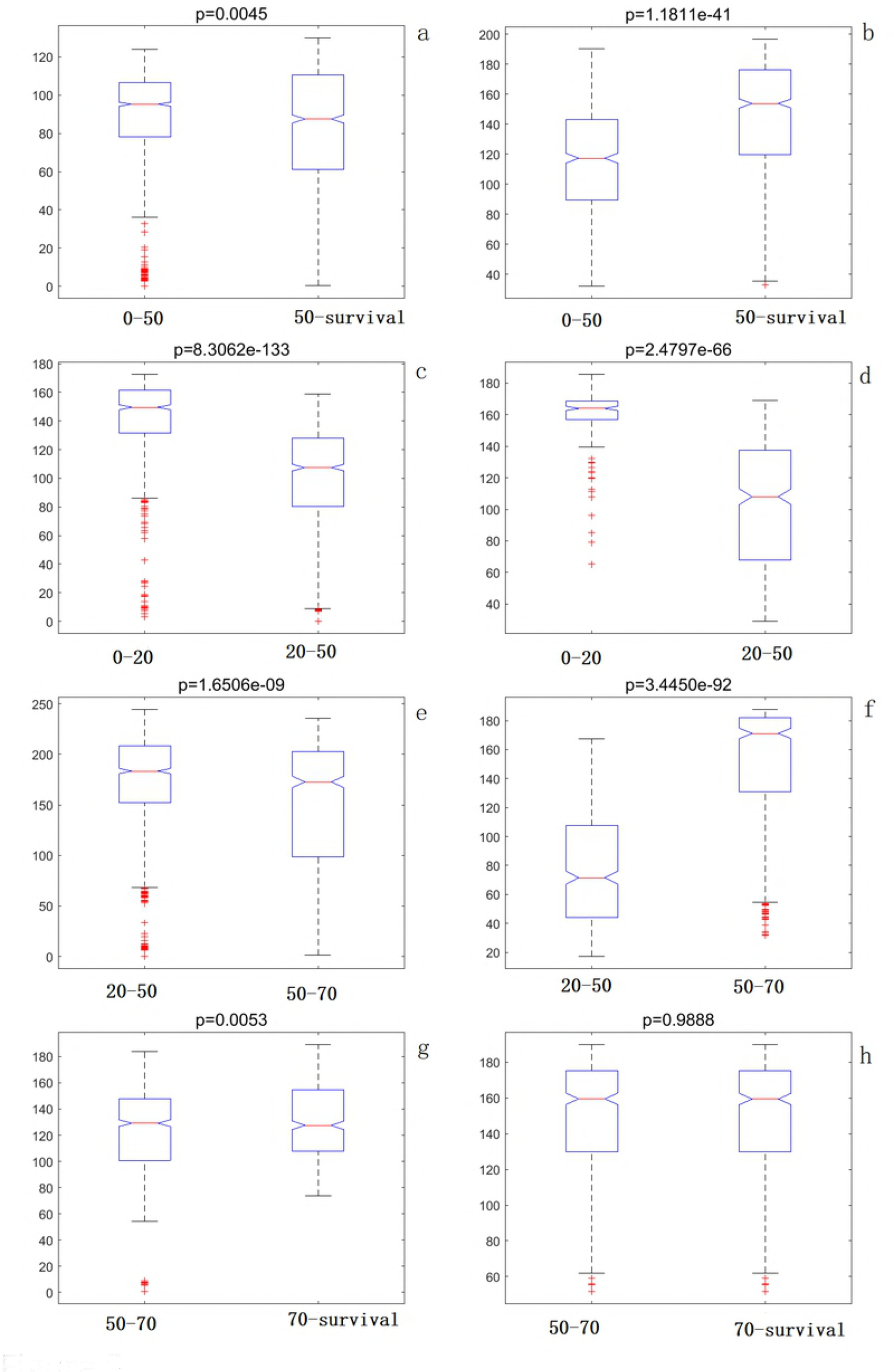
Order-disorder patterns in different aging stages. (a, c, e, g) methylation profiles; (b, d, f, h) expression profiles;

### Aging acceleration of cancers

To study the crosstalk between aging and cancer, the aging associated accelerations were investigated in cancer samples (from the TCGA platform). The order parameters were extracted using the cancer profiles based on each self-organization model, and the module patterns in each model were summarized. Then the aging score were calculated by the module patterns (adjacent normal samples were as the training data, cancer samples were as the test data, and the 0-1 SVM regression was used as the predictor). Strikingly, the results showed that the scores in cancer samples were significantly higher compared to adjacent normal samples, based on both methylation and expression profiles (Figure 6). These results were consistent with previous results, which might reflect the protection of the cancer tissues [5, 17].

**Figure 6.**
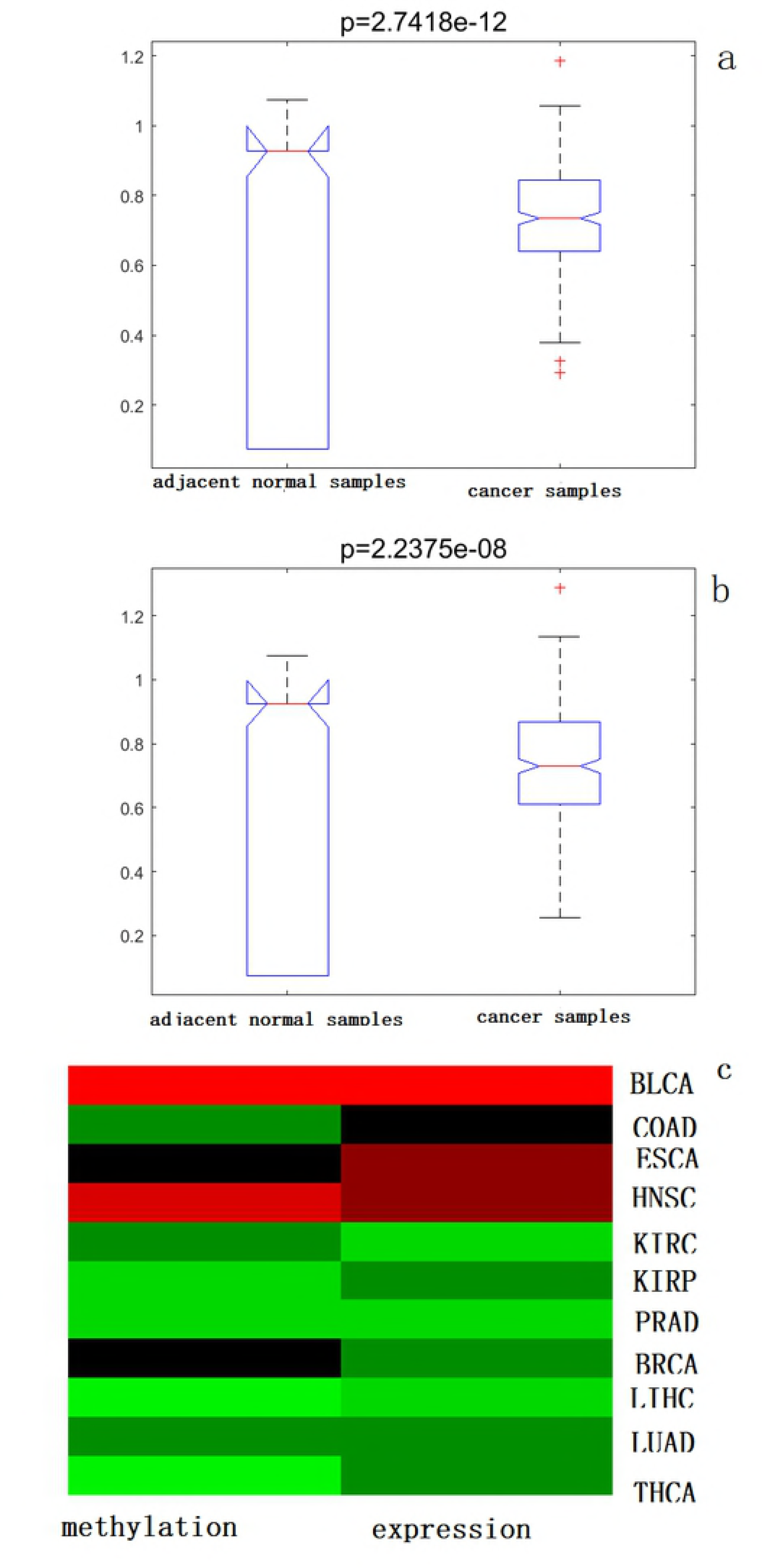
Aging acceleration of cancers. (a) methylation profiles; (b) expression profiles; (c) The heatmap of aging acceleration patterns across cancers;

The correlation between somatic mutation and aging acceleration was also investigate, but none of SNPs with significant (p-value<0.05 and fdr<0.2, using the non-parameter Kruskal-Wallis Test [18]) aging acceleration were identified. The results were perhaps because of not enough samples of paired profiles, aging acceleration tissues in cancer with fewer somatic mutations, or the complexity of the self-organization system [5]. It have been found that there were negative correlation between age acceleration and number of somatic mutations[5]. Therefore, our work also found the negative correlations in most types of cancers (Figure S2 and S3). However, only a few cancers were with significant correlation (e.g. THCA, shown in Figure S2l and S3l).

Further, the 11 types of cancer profiles were clustered based on the mean value of aging acceleration patterns using mean acceleration ratios base on simplified methylation and expression models. As a result (Figure 6c and S4), 7 cancers were identified as one aging acceleration pattern, including BLCA, COAD, ESCA, HNSC, KIRC, KIRP and PRAD; and other 4 cancers were identified as another aging acceleration pattern (BRCA, LIHC, LUAD and THCA). In addition, 49 significant modules were identified based on both differential methylation and expression profiles, where the top differential module (the 926th expression module) connected 50 key BP terms (i.e. cellular localization, skeletal muscle adaptation and GABAergic neuron differentiation) and 51 KEGG pathways (Long-term potentiation and Retinol metabolism), indicating key functions of the aging process of the neuron system. The aging acceleration patterns also revealed basal characteristics across cancers.

## Conclusions

The aging process is regulated by a serious of key markers. The aging markers interact with each other, and performed their functions in the aging specific networks. As a result, identifying modules clustered by these markers during aging was more informative to research the aging process than only finding isolated markers. In this paper, we presented a computational pipeline to model the aging self-organization system and select the order parameters using a serious of supervised learning technologies. The discrimination results showed that our prediction ability was with more accuracy than traditional gene selection / classification methods.

Tissues are usually with declined functions during aging. However, the aging markers interact with each other, and synergistically regulate the aging process. Therefore, the aging process is affected by both ordered and disordered factors. In this work, the complexity characteristics across modules were also found critical patterns in different age groups.

In the immunity theories of aging [19], tissues are involved with progressively functional declines during aging. The functional analysis found the order parameters were enriched in particular BP terms / KEGG pathways in different age groups, where immunity dysfunction and cancer related pathways indicated the common theme of aging. Therefore, the aging process could be predictive by modelling the self-organization system.

Further, the cancer profiles also identified the aging acceleration of cancer samples with statistically significant using the aging scores, based on the aging self-organization system. Both methylation and expression profiles found the cancer samples with aging acceleration compared to normal samples. The results indicated the protective roles of aging in cancers [5]. Moreover, different aging acceleration pattern could also discriminate cancer types.

In summary, we presented the self-organization model of the aging process based on both methylation and expression profiles in this work, where both ordered and disordered critical patterns were identified in different aging stages. Biological features of the order parameters indicated dysfunction of the immunity system and other key functions of aging (i.e. cancers). Thus the aging acceleration also revealed the cross-talk between aging and cancers. In conclusion, the aging self-organization system described here is informative to both aging and aging related diseases.

## Materials and methods

### Data and pre-processing

We obtained methylation and expression profiles from MuTHER study [20] and GEO database (https://www.ncbi.nlm.nih.gov/geo/) with the chronological age (Table S1 and S2), respectively. Only profiles in normal tissues of healthy persons were considered for modelling the aging self-organization systems (samples from normal tissues from persons with cancer and disease status sample / blood, e.g. traumatic blood from healthy persons were discarded). As a result, there were 2226 samples of Gene expression data from 37 datasets, and 4428 samples methylation data from 35 datasets were selected to model the aging self-organization systems, respectively.

For each methylation / expression dataset, the data were treated by a Singular Value Decomposition (SVD) method [16, 21] (regress the first 3 principle components) to assess the sources of inter-sample variation separately in each tissue, and then were normalized to have zero mean and unit variance. Finally the profiles were discretized using two thresholds mean+/-std. If the data came from different platform (e.g. GPL96 / GPL97) even in the same GEO Series, or came from different region of brain (e.g. hippocampus, Posterior cingulate region and so on), the data were treated as independent dataset.

The age groups were partition as: 0-20, 20-50, 50-70 and 70-survival. The choice of age groups was guided by the following criterion: first, the partition of methylation and expression data should be accordance for further integration; second, the human methylation “age acceleration” is significant before age of 20 [5]; third, the sample imbalances between age groups need to be small. Data from different age nearby were used to model the aging self-organization system (0-20 vs. 20-50; 20-50 vs. 50-70; 50-70 vs. 70-sruvival) based on methylation and expression profiles, respectively. Further, simplified classification models were also constructed to discriminate “young” (0-50) and “old” (50-survival) age group [6] based on expression and methylation profiles, respectively.

We also downloaded paired methylation, expression, somatic mutation profiles and clinical data (both cancer and adjacent normal tissue) from the TCGA platform (through the xena website: https://xenabrowser.net/hub/) to further analyze aging related genomic alterations (totally 333 paired samples were obtained).

### The computational pipeline of modelling aging self-organization systems

#### Step 1 filtering the aging background network by maximum mutual information minimum redundancy criterion

Aging is a gradual process with biology functional decline / disorder. The degree of disorder is often evaluated by entropy. The aging process is usually followed energy disperse / entropy increasing; however, in the biological non-closed molecular system, particular bio-markers perform special function of the aging process. Therefore, key order-disorder transitions (or vice versa) might be considered important changes between age group in the aging process.

In this work, the mutual information between different age groups was used to evaluate relevance between genes (i.e. methylation or expression profiles), and the mutual information between genes (from all training samples / age groups) was used to evaluate gene redundancy. In addition, the background interaction system / network of the aging process needs to be satisfied maximum mutual information and minimum redundancy criterion:

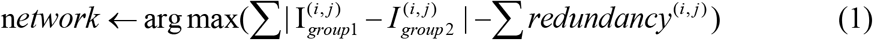

where *I^(i,j)^_group_* indicates the mutual information between genes within the same age group, evaluating the changed dispersion of gene interaction between age groups by the absolute difference, and *redundancy^(i,j)^* indicates the redundancy between genes. As a result, the mutual information between age groups was filtered by the redundancy in the background network as the preliminary gene interaction system of the aging process.

#### Step 2, summarizing gene interactions by the convolution technology

Each “edge” in the aging background network indicating the changed dispersion of gene interaction. As a result, the entire interaction of a gene could be calculated by convoluting all the edges in the background network (i.e. Figure 1b), where the absolute differences of mutual information between age groups were used as the convolution kernel.

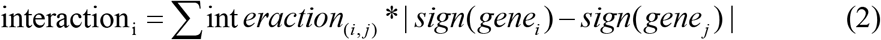

where *gene_i_* and *gene_j_* was the profile of i-th and j-th gene, respectively; and

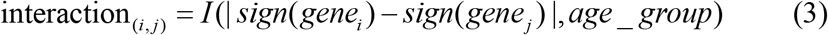

where I(x,y) was the mutual information between x and y.

#### Step 3,calculating the entire pattern by accumulating genes within module

Highly interconnected genes in the network are usually involved in the same biological functions. In this work, genes in the same module were accumulated, where the weights were calculated by the relieff algorithm. Either gene original profiles or convoluted interactions were accumulated was determined by their relieff weights. As a result, each module could be a feature in classification of different age groups. In this work, the SVM classifier (with the linear kernel) was used to discriminate the age groups.

#### Step 4, determining module size by clustering method and cross-validation

The size of module (how many genes in a module) was determined the hierarchical clustering method, where correlation of genes was evaluated by mutual information in the background network between different age groups in the aging process. The clustering degree / times was determined by (5-fold) cross validation.

#### Step 5, identifying order parameters by network sparsification

In this work, only a small ratio of interactions were convoluted in *step 2*, sorted by the mutual information; and only genes with top relieff values were accumulated in the module in *step 2*. sqrt(n) interactions / genes were selected as the order parameters of the aging process using the network sparsification method, where n was the total number interacted with each gene / within the module, respectively.

### Enrichment analysis

Enrichment analyses were carried out to gain significantly biological functions. GO Biological Processes (BP) terms of Gene Ontology (GO) and KEGG pathways were downloaded from Gene Set Enrichment Analysis (GSEA) platform (version 6.1) [22]. The hypergeometric test [23] was performed to estimate the enrichment of these selected genes compared to known GO terms or pathways. Finally, the selected significantly enrichment p-values were controlled by False Discover Rate [24]. The thresholds were set as p-value<0.05 and FDR<0.2.

To evaluate annotated functions across, enriched BP terms / KEGG pathways were calculated by summarizing values of 1-fdr, where fdr<0.2 was set as the threshold.

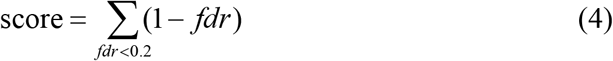

## Supplemental Files

**Figure S1** Hierarchies of the the aging self-organization system.

(a, b) cross-talks between the 492th module and other modules in the model of 0-20 vs. 20-50; (c, d) cross-talks between the 1799th module and other modules in the model of 20-50 vs. 50-70; (a, c) enriched BP terms; (b, d) enriched KEGG pathways;

**Figure S2** Age acceleration versus number of somatic mutations in the TCGA data based on methylation profiles

**Figure S3** Age acceleration versus number of somatic mutations in the TCGA data based on methylation profiles

**Figure S4** aging acceleration characteristics across cancers using the top differential expression module.

(a) connection of BP terms based on order-parameter modules; (b) connection of KEGG pathways based on order-parameter modules;

**Table S1** DNA methylation data involving normal tissues from healthy persons

**Table S2** gene expression data involving normal tissues from healthy persons

**Table S3** modules based on order-parameters of the aging self-organization system using methylation profiles.

**Table S4** modules based on order-parameters of the aging self-organization system using expression profiles.

Note: 0 indicates genes are not selected as order-parameters, otherwise are selected within modules (Table S3-S4).

